# GENESPACE: syntenic pan-genome annotations for eukaryotes

**DOI:** 10.1101/2022.03.09.483468

**Authors:** John T. Lovell, Avinash Sreedasyam, M. Eric Schranz, Melissa A. Wilson, Joseph W. Carlson, Alex Harkess, David Emms, David Goodstein, Jeremy Schmutz

## Abstract

The development of multiple high-quality reference genome sequences in many taxonomic groups has yielded a high-resolution view of the patterns and processes of molecular evolution. Nonetheless, leveraging information across multiple reference haplotypes remains a significant challenge in nearly all eukaryotic systems. These challenges range from studying the evolution of chromosome structure, to finding candidate genes for quantitative trait loci, to testing hypotheses about speciation and adaptation in nature. Here, we address these challenges through the concept of a pan-genome annotation, where conserved gene order is used to restrict gene families and define the expected physical position of all genes that share a common ancestor among multiple genome annotations. By leveraging pan-genome annotations and exploring the underlying syntenic relationships among genomes, we dissect presence-absence and structural variation at four levels of biological organization: among three tetraploid cotton species, across 300 million years of vertebrate sex chromosome evolution, across the diversity of the Poaceae (grass) plant family, and among 26 maize cultivars. The methods to build and visualize syntenic pan-genome annotations in the GENESPACE R package offer a significant addition to existing gene family and synteny programs, especially in polyploid, outbred and other complex genomes.

## INTRODUCTION

*De novo* genome assemblies and gene model annotations represent increasingly common resources that describe the sequence and putative functions of protein coding and intergenic regions within a single genotype. Evolutionary relationships among these DNA sequences are the foundation of many molecular tools in modern medical, breeding and evolutionary biology research. Perhaps the most crucial inference to make when comparing genomes revolves around homologous genes, which share an evolutionary common ancestor and ensuing sequence or protein structure similarity. Analyses of homologs, including comparative gene expression, epigenetics, and sequence evolution, require the distinction between orthologs which arise from speciation events, and paralogs, which arise from sequence duplications. In some systems, this is a simple task where most genes are single copy, and orthologs are synonymous with reciprocal best-scoring BLAST hits. Other sequence similarity approaches such as OrthoFinder (*1*, *2*) leverage graphs and gene trees to test for orthology, permitting more robust analyses in systems with gene copy number (CNV) or presence-absence variation (PAV). However, whole-genome duplications (WGDs), chromosomal deletions, and variable rates of sequence evolution, such as sub-genome dominance in polyploids, can confound the evidence of orthology from sequence similarity alone.

The physical position of homologs offers a second line of evidence that can help to overcome challenges posed by WGDs, tandem arrays, heterozygous-duplicated regions, and other genomic complexities (*3*–*5*). Synteny, or the conserved order of DNA sequences among chromosomes that share a common ancestor, is a typical feature of eukaryotic genomes. In some taxa, synteny is preserved across hundreds of millions of years of evolution and is retained over multiple whole genome duplications (*6*–*8*). Such signals of evolutionary coalescence are often lost in DNA sequences of protein coding genes. Like chromosomal scale synteny, conserved gene order collinearity along local regions of chromosomes can provide evidence of homology, and in some cases enable determinations of whether two regions diverged as a result speciation or a large scale duplication event (*5*). Combined, evidence of gene collinearity and sequence similarity should improve the ability to classify paralogous and orthologous relationships beyond either approach in isolation.

Integrating synteny and collinearity into comparative genomics pipelines also physically anchors the positions of related gene sequences onto the assemblies of each genome. For example, by exploring only syntenic orthologs it is possible to examine all putatively functional variants within a genomic region of interest, even those that are absent in the focal reference genome (*9*). Such a pan-genome annotation framework (*10*) would permit easy access to multi-genome networks of high-confidence orthologs and paralogs, regardless of ploidy or other complicating aspects of genome biology. Here, we present GENESPACE, an analytical pipeline (Supplemental Fig. 1) that explicitly links synteny and sequence similarity to provide high-confidence inference about networks of genes that share a common ancestor, and represents these networks as a pan-genome annotation. We then leverage this framework to explore gene family evolution in flowering plants, mammals and reptiles.

## RESULTS AND DISCUSSION

### GENESPACE methods to compare multiple complex genomes

Until recently, most genome assemblies were haploid, representing meiotically homologous chromosomes as a single haplotype. While this is certainly appropriate for inbred or haploid species, such a representation does not adequately address heterozygosity in outbred species or homeologous chromosomes, which have diverged following a whole-genome duplication in polyploid genomes. With the advent of accurate long-read sequencing, many state-of-the-art genomes of diploid eukaryotes are now phased, representing both homologous chromosomes in the assembly (*10*, *11*). The representation of both meiotically homologous chromosomes in outbred diploids introduces a problem well known in polyploid comparative genomics: paralogs, which are duplicated within a genome, such as homeologs in polyploids or meiotic homologs in outbred diploid genomes, are not as accurately inferred as single-copy orthologs among genomes by graph-based clustering programs. This challenge can be easily addressed in genomes with two complete and easily identifiable sub-genomes (or alternative haplotypes) by splitting chromosomes into separate haploid genomes. However, this splitting approach is not possible in many outbred or polyploid genomes due to chromosomal rearrangements (e.g. maize, see below), and segmental duplications or deletions (e.g. sex chromosomes, see below). Given these known biases, it is crucial to develop a comparative genomics framework that performs adequately in outbred and polyploid genomes.

GENESPACE overcomes the challenge of accurately finding homeologous or meiotically homologous gene pairs by constraining orthogroups (OGs) within syntenic regions. In short, GENESPACE subsets raw global OrthoFinder OGs to synteny by dropping graph edges that span non-syntenic genomic coordinates, thus producing split synteny-constrained OG subgraphs (Supplemental Fig. 1). GENESPACE can then run Orthofinder on BLAST hits within syntenic regions which, when merged with synteny-constrained OGs, produces within-block OGs. Within-block graphs can better capture subgraphs containing distant paralogs because hit scores outside of the focal region are not considered, thereby effectively inferring paralogs with similar efficacy to orthologs (Table 1). GENESPACE then projects the syntenic position of each orthogroup against a single genome assembly of any ploidy, which permits representation of gene presence-absence (PAV) and copy-number (CNV) variation as physically anchored subgraphs along the reference genome. We term this resource a ‘pan-genome annotation’. Since analyses are conducted within syntenic regions, GENESPACE is agnostic to ploidy, duplicated or deleted regions, inversions, or other common chromosomal complexities.

**Table 1.** Summary of orthogroup (‘OG’) inference for polyploids. Orthofinder was run using default settings on three tetraploid inbred cotton genomes (represented as diploid assemblies) and six split sub-genomes. Counts of single-copy orthogroups (more = better) are presented for nine cotton chromosomes.

|  | tetraploid | split by subg. | % split better |
| --- | --- | --- | --- |
| n. *global 1x/homeolog OGs -- | 15,280 | 16,804 | 9.1% |
| n. **synteny-constr. 1x OGs -- | 18,433 | 21,317 | 13.5% |
| n. ***within-block 1x OGs -- | 21,989 | 21,652 | -1.6% |

\*\*Global\* orthogroups were parsed directly from the raw orthofinder (-og) run.
\*\*Synteny-constrained orthogroups are split so that only graph edges within syntenic regions between (sub)genomes are retained. \*\*\*Within block orthogroups are re-calculated from BLAST hits within pairwise syntenic regions.

As a proof of concept, we compared the GENESPACE synteny-constrained orthology inference method with global and sub-genome split OrthoFinder runs using three allotetraploid cotton genomes (*12*). These genomes offer an ideal system to test orthology inference methods due to their easily identifiable sub-genomes, which resulted from an ancient 1.0-1.6 million (M) year ago (ya) whole-genome duplication (WGD), and significant molecular divergence among genomes (160-630k ya). To determine the sensitivity of each approach, we calculated the percent of genes or tandem array representatives captured in orthogroups that were placed in exactly one syntenic position on each sub-genome (Supplemental Fig. 2). Given the known high degree of sequence conservation and little gene presence-absence variation among these cotton genomes and sub-genomes (*12*), most orthogroups should have six syntenic positions across the three cotton genomes, each with two sub-genomes. Therefore, the most accurate method should produce more single-copy orthogroups with exactly six syntenic positions. Given this metric, the run where the sub-genomes were split into separate “species” outperformed the tetraploid run, recovering 9% more orthogroups present only on homologous or homeologous chromosomes across all six sub-genomes. However, GENESPACE’s method to re-run OrthoFinder on synteny constrained within-block BLAST hits effectively brought genome-wide single-copy orthogroup inference in line with the sub-genome split methods (Table 1). These results indicate that, in contrast to previous approaches, GENESPACE infers homeologs between polyploid sub-genomes with similar precision as orthologs among haploid genomes.

In addition to improved accuracy and precision of syntenic orthogroup inference, GENESPACE’s method to find syntenic regions and blocks outperforms collinearity estimates from the program MCScanX (*4*), which serves as an important tool for synteny inference (Table 2). To demonstrate this improvement, we contrasted the two sub-genomes of ‘Pima’ cotton (*Gossypium barbadense*). The 1-1.6M ya divergence between these sub-genomes resulted in many minor and several major inversions and translocations (Supplemental Fig. 2), yet the two genomes remain nearly completely intact and single-copy, excluding tandem arrays. Thus, the vast majority of each sub-genome should correspond to exactly one position in the alternative sub-genome. To test the performance of syntenic block calculations, we tabulated the proportion of 10kb genomic intervals in the expected single-copy dosage or likely erroneous (absent or multi-copy) copy number for three different BLAST hit subsets piped into MCScanX and the complete GENESPACE method (Table 2). MCScanX’s sensitivity causes non-orthologous blocks and overlapping block breakpoints to be included at a high rate: 14% of all intervals were multi-copy in the MCScanX run using raw BLAST hits. However, this issue can be partially resolved by subsetting the BLAST hits to those within the same orthogroups (2.6% multi-copy). This orthogroup constraint performance improvement is the major motivator for the GENESPACE synteny pipeline, which uses orthogroup-constrained BLAST hits as the initial seed for syntenic blocks, then searches all hits within a fixed radius to these anchors. This second proximity search step also resulted in significant gains in single-copy syntenic regions between sub-genomes, simultaneously reducing the amount of un-represented (6.1% to 5.6%) and multi-copy (2.6% to 0.6%) sequences. Combined, these results demonstrate a marked improvement in synteny discovery and block coordinate assignment.

**Table 2.**
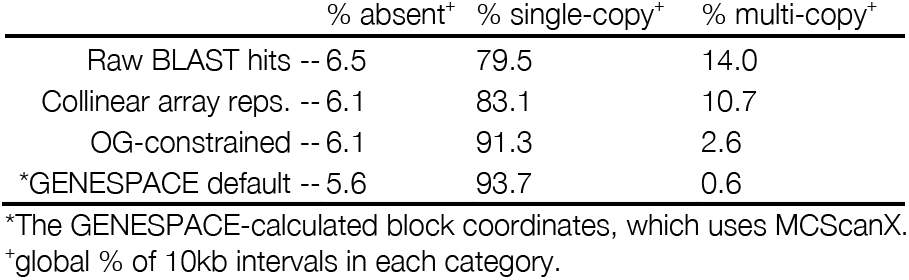
Summary of syntenic blocks between *G. barbadense* sub-genomes. MCScanX_h was run for each subset of BLAST hits and the copy number of each non-overlapping 10kb genomic interval was tabulated from the start/end coordinates of the unique blocks from the collinearity file. The percent of 10kb intervals that are never found within a block (absent), found within exactly one block (single-copy) or in more than one block (multi-copy) are reported.

### Synteny-anchored vertebrate sex chromosomes pan-genome annotations

The GENESPACE pan-genome annotation facilitates the exploration and analysis of sequence evolution across multiple genomes within regions of interest (ROI). Some common use applications include the analysis of QTL intervals (see the next section), or tests of genome evolution at larger phylogenetic scales. One particularly instructive example comes from the origin and evolution of the mammalian XY and avian ZW sex chromosome systems. To explore these chromosomes, we ran GENESPACE on 15 haploid avian and mammalian genome assemblies (Table 3), spanning most major clades of birds, placental mammals, monotremes and marsupials with chromosome-scale annotated reference genomes (Supplemental Fig. 3, Supplemental Data 1-2). We also included two reptile genomes as outgroups to the avian genomes. The heteromorphic chromosomes (Y and W) are often un-assembled, or, where assemblies exist, lack sufficient synteny to provide a useful metric for comparative genomics. As such, we chose to focus on the homomorphic X and Z chromosomes, which have remained surprisingly intact over the >100M years of independent mammalian (*13*) and avian evolution (*14*) (Fig. 1).

**Fig. 1.**
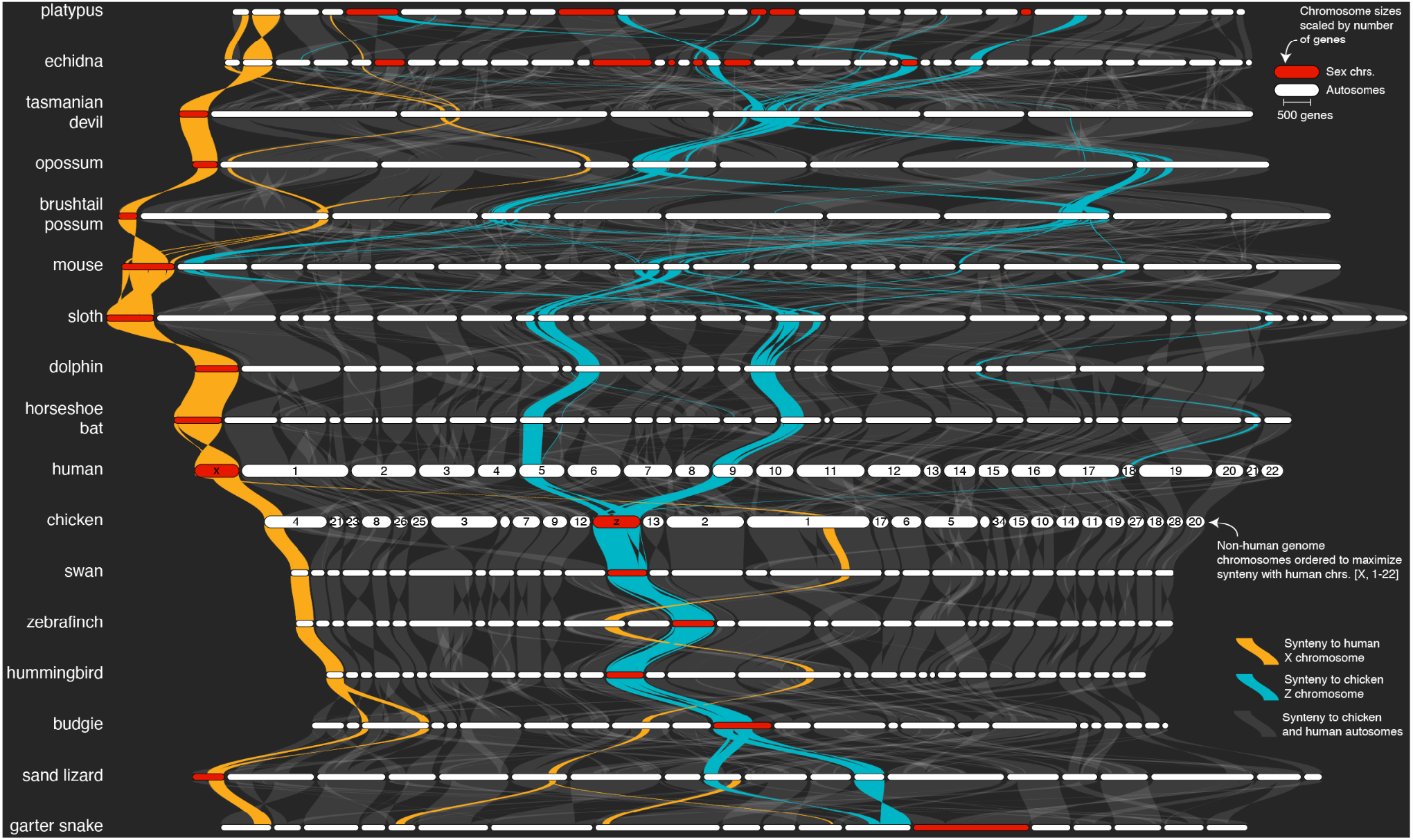
Structural evolution of mammalian X and avian Z sex chromosomes. The reptilian, avian, and mammalian sex chromosomes syntenic network across 17 representative vertebrate genomes (two reptile, five eutherian mammal, three marsupial, two monotremes, and five avian genomes; see Supplemental Fig. 3 for the full synteny graph including autosomes and chromosome labels). The plot was generated by the GENESPACE function plot_riparian. Genomes are ordered vertically to maximize synteny between sequential pairwise genomes. Chromosomes are ordered horizontally to maximize synteny with the human chromosomes [X, Y, 1-22]. Regions containing syntenic orthogroup members to the mammalian X (gold) or avian Z (blue) chromosomes are highlighted. All sex chromosomes are represented by red segments (except the bat chr1, which is most likely the X chromosome but is not represented as such in the assembly), while autosomes are white. Chromosomes are scaled by the total number of genes in syntenic networks and positions of the braids are the gene order along the chromosome sequence.

**Table 3.** Raw data sources. A list of the genomes used in analyses here. Genome version IDs are taken from those posted on the respective data sources and may not reflect the name of the genome in the publication. Where multiple haplotypes are available, only the primary was used for these analyses. All polyploids presented here have only a primary haplotype assembled into chromosomes.

| ID | Species | Genome version | Data source | Ploidy* | Reference |
| --- | --- | --- | --- | --- | --- |
| garterSnake | *Thamnophis elegans* | rThaEle1.pri | NCBI | 1 | (11) |
| sandLizard | *Lacerta agilis* | rLacAgi1.pri | NCBI | 1 | (11) |
| chicken | *Gallus gallus* | mat.broiler.GRCg7b | NCBI | 1 | https://www.ncbi.nlm.nih.gov/grc |
| hummingbird | *Calypte anna* | bCalAnn1_v1.p | NCBI | 1 | (11) |
| budgie | *Melopsittacus undulatus* | bMelUnd1.mat.Z | NCBI | 1 | Unpublished VGP |
| swan | *Cygnus olor* | bCygOlo1.pri.v2 | NCBI | 1 | (11) |
| zebraFinch | *Taeniopygia guttata* | bTaeGut1.4.pri | NCBI | 1 | (11) |
| echidna | *Tachyglossus aculeatus* | mTacAcu1.pri | NCBI | 1 | (34) |
| platypus | *Ornithorhynchus anatinus* | mOrnAna1.pri.v4 | NCBI | 1 | (34) |
| brushtailPossum | *Trichosurus vulpecula* | mmTriVul1.pri | NCBI | 1 | (11) |
| opossum | *Monodelphis domestica* | MonDom5 | NCBI | 1 | (35) |
| tasmanianDevil | *Sarcophilus harrisii* | mSarHar1.11 | NCBI | 1 | (11) |
| human | *Homo sapiens* | GRCh38.p13 | NCBI | 1 | https://www.ncbi.nlm.nih.gov/grc |
| mouse | *Mus musculus* | GRCm39 | NCBI | 1 | https://www.ncbi.nlm.nih.gov/grc |
| dog | *Canis lupus familiaris* | Dog10K_Boxer_Tasha | NCBI | 1 | (36) |
| sloth | *Choloepus didactylus* | mChoDid1.pri | NCBI | 1 | (11) |
| horseshoeBat | *Rhinolophus ferrumequinum* | mRhiFer1_v1.p | NCBI | 1 | (11) |
| dolphin | *Tursiops truncatus* | mTurTru1.mat.Y | NCBI | 1 | Unpublished VGP |
| Phallii | *Panicum hallii* var. *hallii* | HAL2_v2.1 | Phytozome | 1 | (9) |
| switchgrass | *Panicum virgatum* | AP13_v5.1 | Phytozome | 2 | (19) |
| Sviridis | *Setaria viridis* | v2.1 | Phytozome | 1 | (37) |
| Sorghum | *Sorghum bicolor* | BTx623_v3.1 | Phytozome | 1 | (38) |
| maize | *Zea mays* | B73_refgen_v5 | NCBI | *2 | (24) |
| rice | *Oryza sativa* cv 'kitaake' | kitaake_v2.1 | Phytozome | 1 | (39) |
| brachy | *Brachypodium distachyon* | Bd21_v3.1 | Phytozome | 1 | (40) |
| wheat | *Triticum aestivum* | V4 (Chinese Spring) | NCBI | 3 | (41) |
| Gbarbadense | *Gossypium barbadense* | v1.1 | Phytozome | 2 | (12) |
| Gdarwinii | *Gossypium darwinii* | v1.1 | Phytozome | 2 | (12) |
| Gtomentosum | *Gossypium tomentosum* | v1.1 | Phytozome | 2 | (12) |
| 26 NAM parents | *Zea mays* | see data on NCBI | NCBI | *1 | (24) |

\*Ploidy indicates how the genome was treated in the analyses. All values match the ploidy of the primary assembly haplotype except maize, where the refgen\_v5 was treated as diploid (to match both homeologs) in the multi-species run, but as haploid in the NAM founder population to track only meiotic homologs across the population. This parameterization is to match the phylogenetic position of the WGD in the terminal branch of the grass-wide analysis, but ancestral in the 26-NAM analysis.

While the same or similar genomic regions often recurrently evolve into sex chromosomes, perhaps due to ancestral gene functions involved in gonadogenesis, evidence about the non-randomness of sex chromosome evolution is still contentious (*15*). Given our analysis, it is clear that the avian Z chromosome did not evolve from either of the two reptile Z chromosomes sampled here, but instead likely arose from autosomal regions or unsampled ancestral sex chromosomes. The situation in mammals is less clear, in part because both reptile genomes are more closely related to avian than mammalian genomes, which makes ancestral state reconstructions between the two groups less accurate. Nonetheless, the mammalian X and sand lizard Z chromosomes partially share syntenic orthology, an outcome that would be consistent with common descent from a shared ancestral sex chromosome or autosome containing sex-related genes. The shared 91.7M bp region between the human X and sand lizard Z represents 59.0% of the human X chromosome genic sequence. The remaining 64.0M bp of human X linked sequence are syntenic with autosomes 4 (9.9M bp) and 16 (119.6M bp) in sand lizard. The same region is syntenic across three autosomes in the garter snake genome (Fig. 1, Supplemental Data 3).

The eutherian mammalian X chromosome is largely composed of two regions, an X-conserved ancestral sex chromosome region that arose in the common ancestor of therian mammals, and an X-added region that arose in the common ancestor of eutherians (*16*). Consistent with this evolutionary history, the X chromosome is syntenic across all five eutherian mammals studied here. Further, a 107.2M bp (68.8%) segment of the human X, which corresponds with the X-conserved region, is syntenic with 77.8M bp (93.9%) of the tasmanian devil X chromosome and represents the entire syntenic region between the human and all three marsupial X chromosomes (Fig. 1).

Similarly, the chicken Z chromosome is retained in its entirety across all five avian genomes. The only notable exception being the budgie Z chromosome, which also features a partial fusion between the Z and an otherwise autosomal 19.5M bp segment of chicken chromosome 11 (Fig. 1, Supplemental Data 3), potentially representing a neo-sex chromosome fusion that has not yet been described.

In contrast to conserved eutherian and avian sex chromosomes, the complex monotreme X_n_Y_n_ sex chromosomes are only partially syntenic between the two sampled genomes. Only the first X chromosomes are ancestral to both echidna and platypus (*17*), and all are unrelated to the mammalian X chromosomes (Fig. 1, Supplemental Fig. 3), consistent with their independent evolution (*17*). Interestingly, the entirety of the echidna X4 and 47.6M bp (67.9%) of the genic region of the platypus X5 chromosomes are syntenic with the avian Z chromosome (Fig. 1). The phylogenetic scale of the genomes presented here precludes evolutionary inference about the origin of these shared sex chromosome sequences; however, the possibility of parallel evolution of sex chromosomes between such diverged lineages may prove an interesting future line of inquiry.

### Exploiting synteny to track candidate genes in grasses

The Poaceae grass plant family is one of the best studied lineages of all multicellular eukaryotes and includes experimental model species (*Brachypodium distachyon*; *Panicum hallii; Setaria viridis*) and many of the most productive (*Zea mays*- maize/corn; *Triticum aestivum* - wheat, *Oryza sativa* - rice) and emerging (*Sorghum bicolor* - sorghum; *P. virgatum* - switchgrass) agricultural crops. Despite the tremendous genetic resources of these and other grasses, genomic comparisons among grasses are difficult, in part because of an ancient polyploid origin (see the next section), and because subsequent whole-genome duplications are a feature of most clades of grasses. For example, maize is an 11.4M ya paleo-polyploid (*18*), allo-tetraploid switchgrass formed 4-6M ya (*19*), and allo-hexaploid bread wheat arose about 8k ya (*20*). In some cases, homeologous gene duplications from polyploidy have generated genetic diversity that can be targeted for crop improvement; however, in other cases the genetic basis of trait variation may be restricted to sequences that arose in a single sub-genome. Thus, it is crucial to contextualize comparative-quantitative genomics searches and explicitly explore only the orthologous or homeologous regions of interest when searching for markers or candidate genes underlying heritable trait variation — a significant challenge in the complex and polyploid grass genomes. To help overcome this challenge and provide tools for grass comparative genomics, we conducted a GENESPACE run and built an interactive viewer hosted on Phytozome (*21*) among genome annotations for the eight grass species listed above. Owing to its use of within-block orthology and synteny constraints, GENESPACE is ideally suited to conduct comparisons across species with diverse polyploidy events. Default parameters produced a largely contiguous map of synteny even across notoriously difficult comparisons like the paleo homeologs between the maize sub-genomes (Fig. 2a, Supplemental Fig. 4, Supplemental Data 4). Furthermore, the sensitive synteny construction pipeline implemented by GENESPACE effectively masks additional paralogous regions like those from the *Rho* duplication that gave rise to all extant grasses (but see below).

**Fig. 2.**
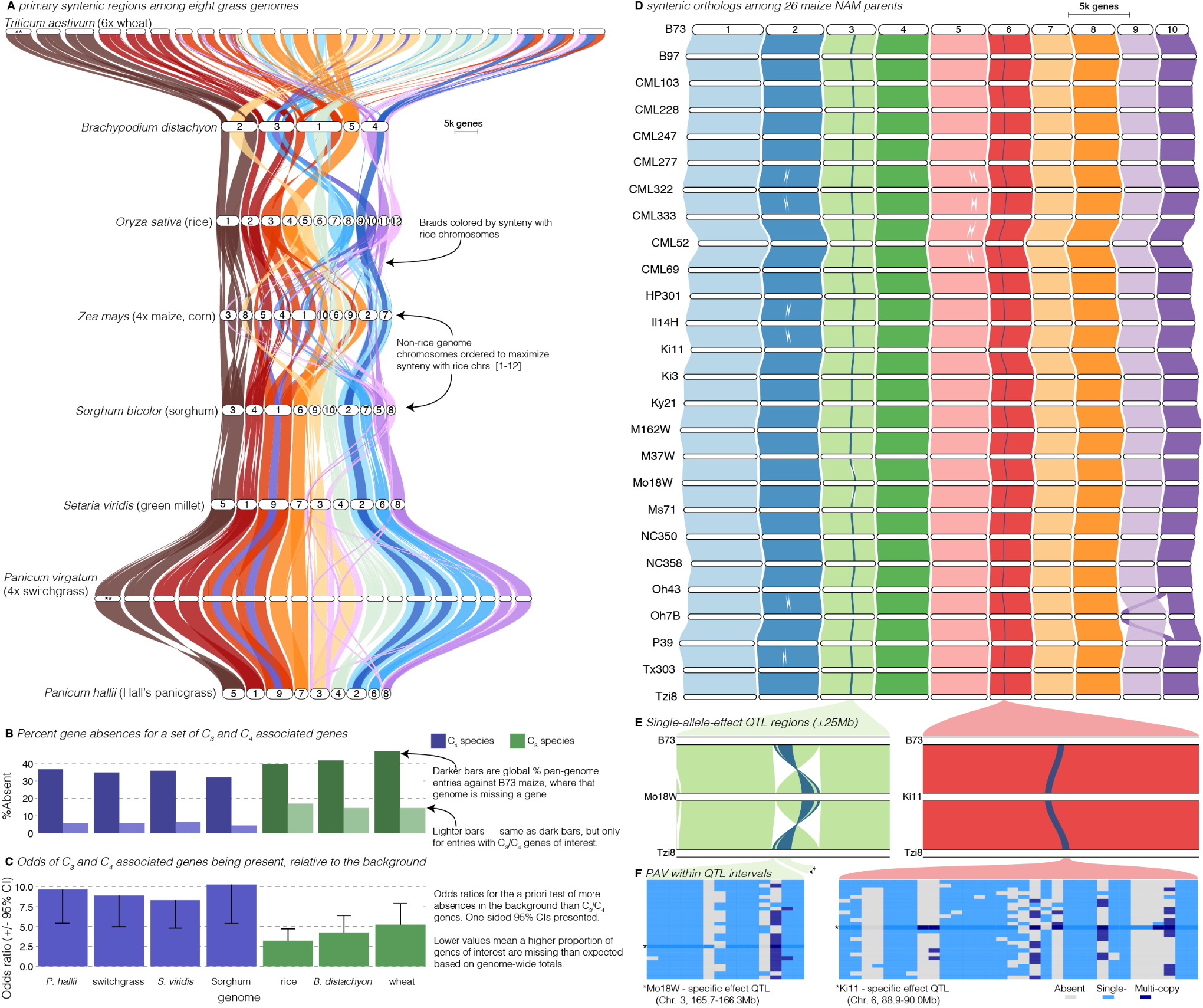
Comparative-quantitative genomics in the grasses. **A** The GENESPACE syntenic map (‘riparian plot’) of orthologous regions among eight grass genomes. Chromosomes are ordered to maximize synteny with rice and ribbons are color-coded by synteny to rice chromosomes. Chromosome names are too long to fit for the neo-polyploids (**); Supplemental Figure 4 contains names of all chromosomes. **B** The upper bars display the proportion of maize gene models without syntenic orthologs (“absent”) in each genome, split by the full background (dark colors) and 86 genes annotated for roles in the evolution of C3/C4 photosynthesis. **C** The proportion of absent genes is higher in the C3 genomes (green bars), even when controlling for more global gene absences (lower odds ratios). **D** Syntenic orthologs, controlling for homeologs among the 26 maize NAM founder genomes, with two general QTL intervals highlighted. **E** Focal QTL regions that affect productivity in drought where only the genome that drives the QTL effect (middle genome); the top (B73) and bottom (Tzi8) genomes are presented and the region plotted is restricted to the 50Mb physical B73 interval surrounding the QTL. Note that the chr3 QTL disarticulates into two intervals. Due to a larger number of potential candidate genes, the larger chr3 region, flagged with **, is explored separately in Supplemental Figure 6. **F** Presence-absence and copy number variation are presented for two of the three intervals. The focal genome is flagged * and its corresponding map colors are more saturated.

Breeders and molecular biologists can take two general approaches to understanding the genetic basis of complex traits: studying variation caused by *a priori*-defined genes of interest, or determining candidate genes from genomic regions of interest. As an example of the exploration of lists of *a priori*-defined candidate genes, we analyzed the functional and presence-absence variation of 86 genes shown to be involved in the transition between C_3_ and C_4_ photosynthesis (*22*), the latter permitting ecological dominance in arid climates and agricultural productivity under forecasted increased heat load of the next century. To conduct this analysis, we built pan-genome annotations across the seven grasses anchored to C_4_ maize (Supplemental Data 5), which was the genome in which these genes were discovered. This resulted in 159 pan-genome entries; nearly always two placements for each gene in the paleo-tetraploid maize genome. Given that many of these genes were discovered in part because of sequence similarity to genes in *Arabidopsis* and other diverged plant species, it is not surprising that PAV among C_3_/C_4_ genes was lower than the background (9.7% vs 38.2%, odds = 5.7, P < 1×10^−16^; Fig. 2b). However, these ratios were highly variable among genomes, particularly among the C_3_ species (wheat, rice, *B. distachyon*), which had far higher percent absences than the C_4_ species (15.3% vs. 5.5%, odds = 3.1, P = 6.25×10^−8^, Fig. 2b). This effect is undoubtedly due in part to the increased evolutionary distance between maize and the C_3_ species compared to the other C_4_ species. However, when controlling for the elevated level of absent genes globally in C_3_ species, the effect was still very strong: the odds of C_3_ species having more of these C_3_/C_4_ genes at syntenic pan-genome positions than the background was always lower than the C_4_ species (Fig. 2c). Despite these interesting patterns, given only a single C_3_/C_4_ phylogenetic split in this dataset, it is impossible to test evolutionary hypotheses regarding the causes of such PAV. Nonetheless, this result suggests a possible role of gene loss or gain as an evolutionary mechanism for drought- and heat-adapted photosynthesis.

Like the exploration of *a priori*-defined sets of genes, finding candidate genes within quantitative trait loci (QTL) intervals usually involves querying a single reference genome and extracting genes with promising annotations or putatively functional polymorphism. In the case of a biparental mapping population genotyped against a single reference, this is a fairly trivial process where genes within physical bounds of a QTL are the candidates. However, many genetic mapping populations now have reference genome sequences for all parents; this offers an opportunity to explore variation among functional alleles and presence absence variation, which would be impossible with a single reference genome. GENESPACE is ideally suited for this type of exploration, and indeed was originally designed to solve this problem between the two *P. hallii* reference genomes and their F_2_ progeny (*9*) using synteny to project the positions of genes across multiple genomes onto the physical positions of a reference.

To illustrate this approach, we re-analyzed QTL generated from the 26-parent USA maize nested association mapping (NAM) population (*23*). Originally, candidates for these QTL were defined by the proximate gene models only in the B73 reference genome (*23*); however, with GENESPACE and the recently released NAM parent genomes (*24*), it is now possible to evaluate candidate genes present in the genomes of other NAM founder lines but either absent or unannotated in the B73 reference genome. We built a single-copy synteny graph of all 26 NAM founders, anchored to the B73 genome to explore this possibility (Fig. 2d; Supplemental Data 6; Supplemental Fig. 5) and extracted the three QTL intervals (Fig. 2d–e) where the allelic effect of a single parental genome was an outlier relative to all other alleles. Such ‘private’ allelic contributions, which may be driven by parent-specific sequence variation, were manifest here as delayed period of silking-anthesis of progeny with the Mo18W allele at two adjacent Chr3 QTLs and reduced plant height under drought for progeny with the Ki11 allele at the Chr6 QTL (*23*). Given that these QTL were chosen only due to their parental allelic effects, we were surprised to find that the two Mo18W QTL regions exist within a 11.7M bp derived inversion that is only found in the Mo18W genome (Fig. 3d–e). Since inversions reduce recombination, it is possible that multiple Mo18W causal variants have been fixed in linkage disequilibrium in this NAM population. In addition to this chromosomal mutation and sequence variation between the parents and B73 (*23*), we sought to define additional candidate genes from the patterns of presence-absence and copy-number variation, explicitly looking for genes that were private to the focal genome. Two genes in the smaller chr3 and one gene in the larger chr3 interval were private to Mo18W and four genes in three pangenome entries (one two-member array) were private to Ki11 in the chr6 interval (Fig. 2f, Supplemental Fig. 6, Supplemental Data 7). While none of these genes have functional annotations relating to drought, this method provides additional candidates that would not have been discovered by B73-only candidate gene exploration.

**Fig. 3.**
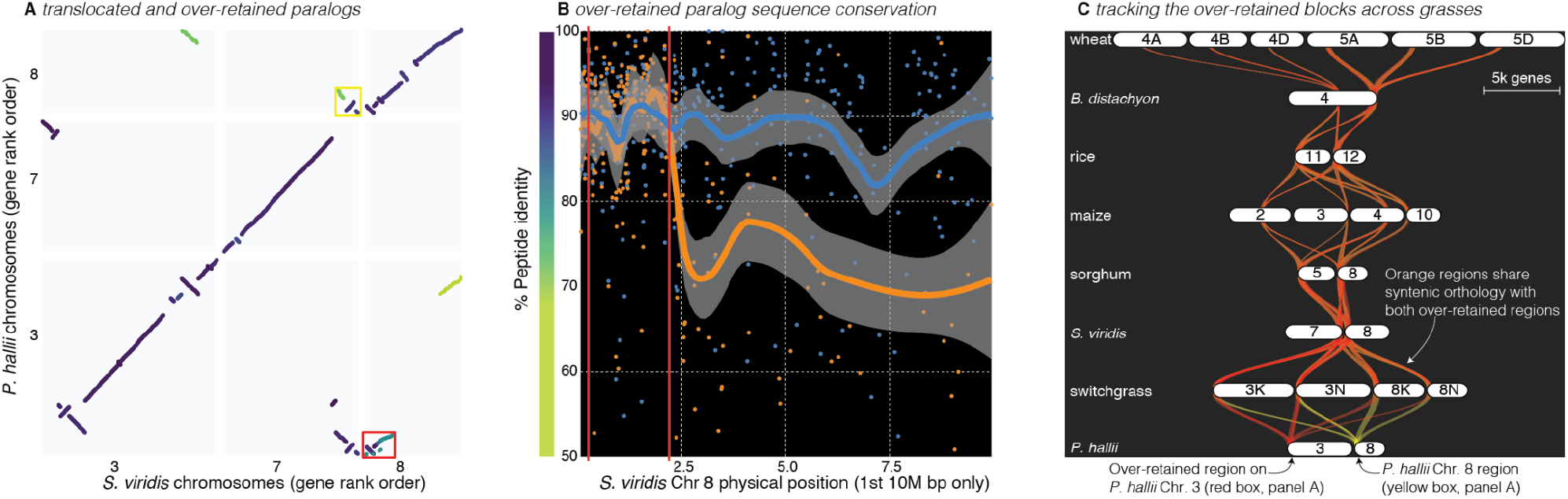
Analysis of the grass *Rho* WGD. **A** Syntenic anchor blast hits where the target and query genes were in the same orthogroup between *P. hallii* and *S. viridis* genomes. The color of each point indicates the peptide identity of each pair of sequences; the color scale is shown along the y axis of panel B . **B** The protein identity of *S. viridis* chromosome 8 primary orthologous (blue line) hits against *P. hallii* chromosome 8 and the secondary hits (orange line) against *P. hallii* chromosome 3 demonstrate sequence conservation heterogeneity. The region between the two red vertical lines corresponds to the red-boxed over-retained primary block in panel A. **C** The two boxed regions in panel A were tracked from their origin on *P. hallii* chromosome 3 (red) and 8 (yellow). Note that all syntenic orthologous regions across the graph contain both P*. hallii* source regions (50% transparency of the braids - overlapping regions appear orange).

### Studying the whole-genome duplication that led to the diversification of the grasses

Like most plant families (*25*–*27*), but unlike nearly all animal lineages (*28*), the grasses radiated following a whole-genome duplication: the ~70M ya *Rho* WGD. The resulting gene family redundancy and gene-function sub-functionalization is hypothesized to underlie the tremendous ecological and morphological diversity of grasses (*29*–*31*). To explore sequence variation among *Rho*-derived paralogs, we used GENESPACE to build a ploidy-aware syntenic pan-genome annotation among these eight species (Supplemental Data 8), using the built-in functionality that allows the user to mask primary (likely orthologous) syntenic regions and search for secondary hits (likely paralogous, Fig. 3a). Overall, the peptide identity between *Rho*-derived paralogous regions was much lower than orthologs among species (e.g. *S. viridis* vs. *P. hallii*: Wilcoxon *W* = 88094632, *P* < 10^−16^; Supplemental Data 9), consistent with the previous discovery that the *Rho* duplication predated the split among most extant grasses by >20M years (*32*). However, as has been previously observed, there is significant variation in the relative similarity of *Rho*-duplicated chromosome pairs (*33*). As an example, the peptide sequences of single-copy gene hits in primary syntenic regions (median identity = 90.6%) between chromosome 8 of *P. hallii* and *S. viridis*, were 26.9% more similar than the secondary *Rho*-derived regions (median identity = 71.4%, Wilcoxon *W* = 87842, *P* < 10^−16^). However, *S. viridis* chromosome 8 contained a single paralogous region between all seven grass genomes that could not be distinguished from the primary regions, based on synteny or orthogroup identity. Unlike all other *Rho*-derived blocks, the *P. hallii* paralogs to this 2.7M bp chromosome 8 region were not significantly less conserved than the primary orthologous region (91.6% vs. 91.9%, *W* = 14830, *P* = 0.13). Outside of this region, the peptide identity of paralogs dropped back to the genome-wide average (Fig. 3b).

Indeed, the GENESPACE run treating the eight genomes as haploid representations could not distinguish between the *Rho* derived paralogs in the over-retained region across all grasses (Fig. 3c), with the exception of all chromosome pairs between *B. distachyon* and wheat and blocks connecting Maize chromosome 10 to sorghum chromosome 5. It is interesting to note that all syntenic over-retained regions are at the extreme terminus of the chromosomes outside of maize, *B. distachyon* and wheat; further, the only genome with complete segregation of the two paralogs, wheat, also retains these regions in the center of all six chromosomes (Fig. 3c). These results are consistent with the proposed evolutionary mechanism (*33*) where concerted evolution and “illegitimate” homeologous recombination may have homogenized these paralogous regions. This process would be less effective in pericentromeric regions than the chromosome tails, where a single crossover event would be sufficient to homogenize two paralogous regions that arose 70M ya.

## Conclusions

Combined, the historical abundance of genetic mapping studies and ongoing proliferation of genome resources provides a strong foundation for the integration of comparative and quantitative genomics to accelerate discoveries in evolutionary biology, medicine, and agriculture. The incorporation of synteny and orthology into comparative genomics and quantitative genetics pipelines offers a mechanism to bridge these disparate disciplines. Here, we presented the GENESPACE R package and the syntenic pan-genome annotation as a framework to help bridge the current gaps between comparative and quantitative genomics, especially in complex evolutionary systems. We hope that the examples presented here will inspire further work to leverage the powerful genome-wide annotations that are coming online, both within and among species.

## METHODS

All analyses were performed in R 4.1.2 on macOS Big Sur 10.16. The following R packages were used either for visualization or within GENESPACE v0.9.3 (11-February 2022 release): data.table v1.14.0 (*42*), dbscan v1.1-8 (*43*), igraph v1.2.6 (*44*), Biostrings v2.58.0 (*45*), rtracklayer v1.50.0 (*46*). GENESPACE also calls the following third party software: diamond v2.0.8.146 (*47*), OrthoFinder v2.5.4 (*1*), and MCScanX no version installed on 10/23/2020 (*4*).

All results, tables (except Table 3), figures (except Fig. S1) and statistics were generated programmatically; the accompanying scripts and key output are available on github: jtlovell/GENESPACE_data. Minor adjustments to figures to improve clarity were accomplished in Adobe Illustrator v26.01. Below, we provide a high-level description of the GENESPACE pipeline and the methods to produce the results presented here. A full description of each step in GENESPACE is provided in the documentation that accompanies the package source code on github (jtlovell/GENESPACE).

### Description of the vignettes

Raw genome annotations were downloaded on or before 8-October 2021. See Table 3 for data sources, citations and metadata. For the analyses presented here, we conducted six GENESPACE runs: cotton tetraploid, cotton sub-genome-split, vertebrates, grasses, grass *Rho* duplication, and maize 26 NAM parents.

All GENESPACE runs used default parameterization, with the following exceptions: (1) both cotton runs used a minimum block size and maximum number of gaps of 10 (default = 5 for both), (2) the *Rho* grass run allowed a single secondary hit (default is 0, this is how the paralogs are explicitly searched for) and maximum number of gaps in secondary regions of 10 (default is 5, relaxed to reduce ancient paralogous block splitting), and (3) the maize run used the “fast” OrthoFinder method since all genomes are closely related and haploid. Some maize genomes contain small alternative haplotype scaffolds, which were dropped for all analyses.

The cotton runs employed the GENESPACE “outgroup” functionality, which allows the user to specify a genome that is included in the seed OrthoFinder run, but is ignored for all synteny and pan-genome construction steps. This can be important when dealing with highly diverged species that do not share complete synteny, but are needed for accurate orthogroup inference. For example, a run with only the three cotton genomes would be likely to split sub-genome orthogroups since the WGD predated speciation. As such, we included *Theobroma cacao* (*48*) as an outgroup.

The publicly available C3/C4 gene lists and QTL intervals were generated against the v2 maize assembly. To make this comparable to the across-grass and NAM parent GENESPACE runs, we also accomplished a fast GENESPACE run between v2 and the two v5 versions used here. The orthologs and syntenic mapping between these versions are included as text files in the data repository.

Statistics presented here were all calculated within R. To compare non-normal distributions (e.g. sequence identity), we used the non-parametric signed Wilcoxon ranked sum test. To measure sequence divergence, we conducted pairwise peptide alignments via Needleman-Wunsch global alignment, implemented in the Biostrings (*45*) function, pairwiseAlignment. We then used this alignment to calculate the percent peptide sequence identity between the un-gapped aligned regions for any two single-copy anchor hits using the Biostrings function pid with the type2 method. To determine single outliers from a unimodal distribution, we applied the Grubbs test implemented in the outliers R package (*49*). Some figures were constructed outside of GENESPACE using base R plotting routines and ggplot2 v3.3.3 (*50*). Some color palettes were chosen with RColorBrewer (*51*) and viridis (*52*).

### GENESPACE pipeline: Running orthofinder within R

GENESPACE operates on gff3-formatted annotation files and accompanying peptide fasta files for primary gene models. There are convenience functions for re-formatting the gff and peptide files to simplify the naming scheme and reduce redundant gene models to the primary longest transcript. With these data in hand OrthoFinder (*1*) is run on the parsed primary peptide files. While the default behavior of GENESPACE is to run OrthoFinder using its default parameters (diamond2 --more-sensitive), GENESPACE also offers a ‘fast’ method that performs only one-way diamond2 (*47*) searches, where the genome annotation with more gene models serves as the query and the smaller annotation is the target. The diamond BLAST-like (hereon ‘BLAST’) results are mirrored and each are stored as OrthoFinder-formatted blast8 text files. OrthoFinder is then run to the orthogroup-formating step (-og) on the pre-computed BLAST text files. This method results in significant speed improvements with little loss of fidelity among closely-related haploid genomes (Table 4).

**Table 4.** Comparison of GENESPACE setting performance. The mirrored ‘fast’ method significantly speeds up orthofinder runs by calling diamond blastp --fast on each non-redundant pairwise combination of genomes. However, this approach is less sensitive than the default performance and is suggested for only closely-related haploid genomes, as the recall of 2:2:2 OGs is slightly less sensitive than the default specification.

|  | Default orthofinder | GENESPACE ‘fast’ |
| --- | --- | --- |
| n. 1:1:1 OGs | 22,050 | 22,444 |
| n. 2:2:2 OGs | 13,793 | 13,511 |
| n. tandem arrays | 10,597 (4433) | 10,599 (4426) |
| *Run time (minutes) | 59.95 | 12.45 |

\*Run time is for ortholog/orthogroup inference, not the GENESPACE pipeline as a whole, using the three unsplit cotton genomes, running on 6 2Gb cores.

There are two methods to infer orthogroups; the original (-og) method clusters genes and builds an undirected cyclic graph from closely related genes bases on BLAST scores (*2*), while hierarchical phylogenetic orthogroups can disarticulate the clustered orthogroups based on gene trees (*1*). The latter approach may more effectively exclude paralogs from orthogroups (Supplemental Table 1). Finally, orthofinder infers pairwise orthologs as directed acyclic graphs from one genome to each other (*1*). The orthologs represent the most strict definition of orthology and are based on gene trees. GENESPACE attempts to merge the benefits of each of these methods by first, only considering -og orthogroups for synteny, which allows users to optionally include paralogs in the scan. If hierarchical orthogroups were used instead, a dramatic decrease in homeologous gene discovery would be expected. To take advantage of the more advanced orthofinder methods, GENESPACE includes non-syntenic gene tree-inferred orthologs into the pan-genome annotation during its final steps (see below).

Orthofinder defines orthogroups as *the set of genes that are descended from a single gene in the last common ancestor of all the species being considered*. As such, the scale of the orthofinder run matters, often significantly. For example, an orthogroup would not be likely to contain homeologs across the two ancient sub-genomes for an orthofinder run that included only two maize genomes — since the coalescence of any two maize genotypes occurred well before the ~12M ya whole genome duplication, few homeologs would both be descended from the same common ancestor when considering only maize genotypes. This is why the within-maize NAM parent run (Fig. 2d) excludes homeologs. However, if an outgroup to maize is included in the orthofinder run, both maize homeologs would be likely to show common ancestry to a single gene in the outgroup, thus connecting the maize homeologs into a single orthogroup. This is why both maize homeologous regions are present in the across-grasses synteny graph (Fig. 2a) despite using identical parameters to the maize NAM parent run. Given the potentially significant role of outgroups on the results of the global orthofinder run (Supplemental Table 1), GENESPACE offers an “outgroup” parameters, which specifies which of the genomes should be included in the orthofinder run, but excluded for all downstream analyses.

### GENESPACE pipeline: Build syntenic orthogroup graphs

Syntenic regions are extracted from BLAST hit files with graph- and cluster-based approaches using a set of user-defined parameters. While these parameters allow for flexibility, the defaults are sufficient for most high-quality genomes and evolutionary scenarios; for example, we used the same default parameters for 300M years of vertebrate evolution, 65M years and multiple WGDs of grasses, and 10k years of Maize divergence. For a full list of parameters, see documentation of the set_syntenyParams GENESPACE function, but here, we will discuss the (1) the minimum number of unique hits within a syntenic block (‘blkSize’, default = 5), (2) the maximum number of gaps within a block alignment (‘nGaps’, default = 5), and (3) the radius around a syntenic anchor for a hit to be considered syntenic (‘synBuff’, default = 100).

Prior to pairwise synteny searches, ‘collinear arrays’ are defined for each genome as groups of genes separated by no more than synBuff genes on the same chromosome that share an orthogroup. For each collinear array, the single physically most central gene is flagged as the ‘array representative’. Only the array representatives can be syntenic anchors (see below); this culling produces more accurate block coordinates in regions with large tandem arrays (Table 2) and substantial speed improvements in highly repetitive genomes.

For each pairwise combination of genomes, synteny is inferred in three steps: (1) the potential syntenic anchor hits are extracted as the top n hits for each array representative gene (where n is the expected ploidy of the alternate genome); (2) collinear anchors are defined by MCScanX; (3) hits within a buffer radius of the collinear anchors are extracted by dbscan. For intra-genomic searches within a haploid genome, synteny is simply defined as the region within the synBuff of self hits. Intra-genomic searches within polyploids (or outbred diploids) are more complicated, as self-hits will cause non-self regions to appear highly broken up. To resolve this issue, the self-hit regions are masked and syntenic regions are calculated on the non-self space following the method for inter-genomic synteny. Syntenic orthogroups, which are initially defined as synteny-constrained global orthogroups, can be updated to include re-calculated within-block orthogroups. This step is computationally intensive and yields significantly improved results only when one or more of the genomes are not haploid (Table 1). As such, the default behavior of GENESPACE is to only run within-block OrthoFinder when any of the genomes have diploid or higher ploidy.

### GENESPACE pipeline: Constructing pan-genome annotations

Pairwise syntenic orthologs are decoded into a multi-genome pan-annotation, which is represented by a text file containing the expected position of all syntenic orthologs across all genomes. This dataset is built in three steps: First, a reference pan-genome annotation is built for all syntenic orthogroups that include a hit in the user-specified reference genome, producing a synteny-aware database that represents each directed subgraph containing a reference genome gene across all genomes. Second, the expected physical position of all genes are interpolated from the syntenic block anchor hits and orthogroups missing from the reference pan-genome annotation are added accordingly, which permits inference of presence-absence variation within a physical position. These interpolated positions are integrated into the pan-genome annotation where each subgraph in the pan-genome is checked as to whether it has a representative anchored in the reference pan-genome. Third, non-syntenic orthologs are extracted from the raw orthofinder run and added to the pan-genome annotation. The reference pan-genome contains all syntenic orthogroup hits connected by a directed acyclic graph to a reference gene. However, there are many cases where the reference gene in this graph is not the only mapping to the reference. For example, polyploids should have multiple positions. As such, we need to cluster the reference positions of all genes in all subgraphs to ensure that all syntenic positions and PAV are captured accurately.

## Acknowledgements

The GENESPACE pipeline has been improved by advice and testing by A. Healey, N. Walden, V. Markham, R. Walstead, S. Carey, L. Smith, J. Vogel, J. Willis, J. Jenkins, T. Juenger and many others. Thanks to J. Schnable, J Leebens-Mack, J.G. Monroe, C.H. Li, R. Tarvin, and M. Hufford for help refining the datasets and analyses presented in this manuscript. Thank you to Erich D. Jarvis and the Vertebrate Genome Project members for advice and pre-publication access to several genomes (budgerigar and dolphin). The work conducted by the US Department of Energy Joint Genome Institute is supported by the Office of Science of the US Department of Energy under Contract No DE-AC02-05CH1123. Visualization was inspired in part by MCScanX and pairwise ‘river’ plots generated by other software. The use of syntenic orthogroups was originally inspired by work developed by CoGe. MAW’s work on this was supported by the National Institute of General Medical Sciences (NIGMS) of the National Institutes of Health (NIH) grant R35GM124827. JTL would like to thank Ashley Lovell, our friends and family for their support, which allowed him to work on this project during the difficult past two years.

## Data availability

Raw data was sourced entirely from NCBI and Phytozome. Processed data, intermediate files, scripts, plots and source data are all available in the data repository: https://github.com/jtlovell/GENESPACE_data. All source code and documentation for the GENESPACE R package can be found at https://github.com/jtlovell/GENESPACE. An interactive viewer for the plant genomes can be found on phytozome at https://phytozome-next.jgi.doe.gov/tools/dotplot/synteny.html.

## Description of supplemental data

**Supplemental Data 1**. Pan-genome annotation of the vertebrates using the human genome as the reference coordinate system. For each row (pan-genome entry), there is position information, projected against the gene order coordinate system of the human genome; pgChr and pgOrd are the human chromosome and gene rank order position of that entry. There is also a pgID column, which splits entries that happen to be at the same position but lack a reference gene. The remaining columns are the 17 vertebrate genome IDs. In each column, syntenic orthogroup (un-flagged), non-syntenic orthologs (flagged *) and tandem array members (flagged +) are ‘|’ separated.

**Supplemental Data 2**. Pan-genome annotation of the vertebrates using the chicken genome gene rank order as the reference coordinate system. Columns follow supplemental data 1.

**Supplemental Data 3**. Physical coordinates of syntenic block breakpoints among all pairwise combinations of the 17 vertebrate genomes. Pairwise combinations are distinguished by the genome IDs presented in the first two columns. The following six columns (chr1, chr2, start1, start2, end1, end2) are separated where columns ending in “1” belong to the coordinate system of the genome ID in the first “genome1” column, while columns ending in “2” belong to the coordinate system of the genome ID in the second “genome2” column. Start and end coordinates are in base pairs. Orientation, column “orient” is flagged as “+” for collinear, “-” for inverted. The last column, “nhits” is the number of syntenic anchor hits within that block.

**Supplemental Data 4**. Physical coordinates of syntenic block breakpoints among all pairwise combinations of the 8 grass genomes. Columns follow supplemental data 3.

**Supplemental Data 5**. Pan-genome annotation of the grasses using the maize B73 genome gene rank order as the reference coordinate system. Columns follow supplemental data 1.

**Supplemental Data 6**. Physical coordinates of syntenic block breakpoints among all pairwise combinations of the 26 NAM parents. Columns follow supplemental data 3.

**Supplemental Data 7**. Pan-genome annotation of the 26 NAM parents using the maize B73 genome gene rank order as the reference coordinate system. Columns follow supplemental data 1.

**Supplemental Data 8**. Pan-genome entries of the 26 maize NAM founders for each of the three QTL regions in Li et al. 2016. Columns follow supplemental data 1, with the additional first column “qtl”, which holds the QTL id, coded as [phenotype] [(private focal genome)]: [chromosome], [start Mbp]-[end Mbp].

**Supplemental Data 9**. Pan-genome annotation of the grasses, explicitly including the *Rho*-duplicated homologs into the graph, and using the *S. viridis* genome as the reference coordinate system. Columns follow supplemental data 1.

**Supplemental Data 10**. Hits between *P. hallii* and *S. viridis* genes that are members of the same within-block orthogroups and are syntenic anchors. The first 12 columns (id1, id2, genome1, genome2, chr1, chr2, start1, end1, ord1, start2, end2, ord2) are separated where columns ending in “1” belong to the coordinate system of the genome ID in the first “genome1” column, while columns ending in “2” belong to the coordinate system of the genome ID in the second “genome2” column. Start and end are bp positions, ord is the gene rank order. The two measures of percent protein identity are given in pid1 and pid2 columns. The block type, categorized as orthologous (“orth”), over-retained *Rho* paralog (“overr”), regular *Rho* paralog (“rho”) and ambiguous (“ambig”) are given in the column blockType.

## SUPPLEMENTAL FIGURES

**Supplemental Figure 1.**
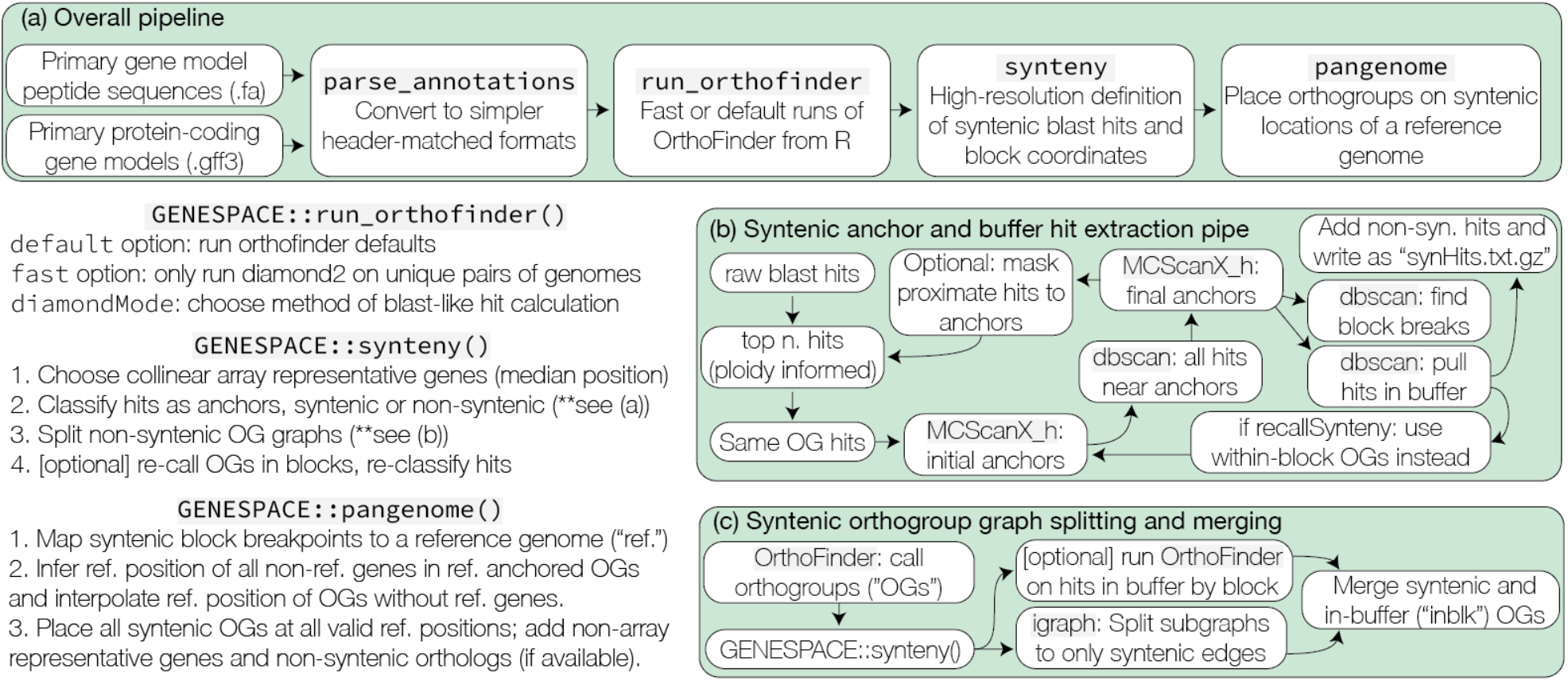
Description of the pipeline. Green boxes show the primary (a), synteny (b) and syntenic orthogroup (c) modules. Verbal descriptions of the three main GENESPACE functions are presented in the bottom right.

**Supplemental Figure 2.**
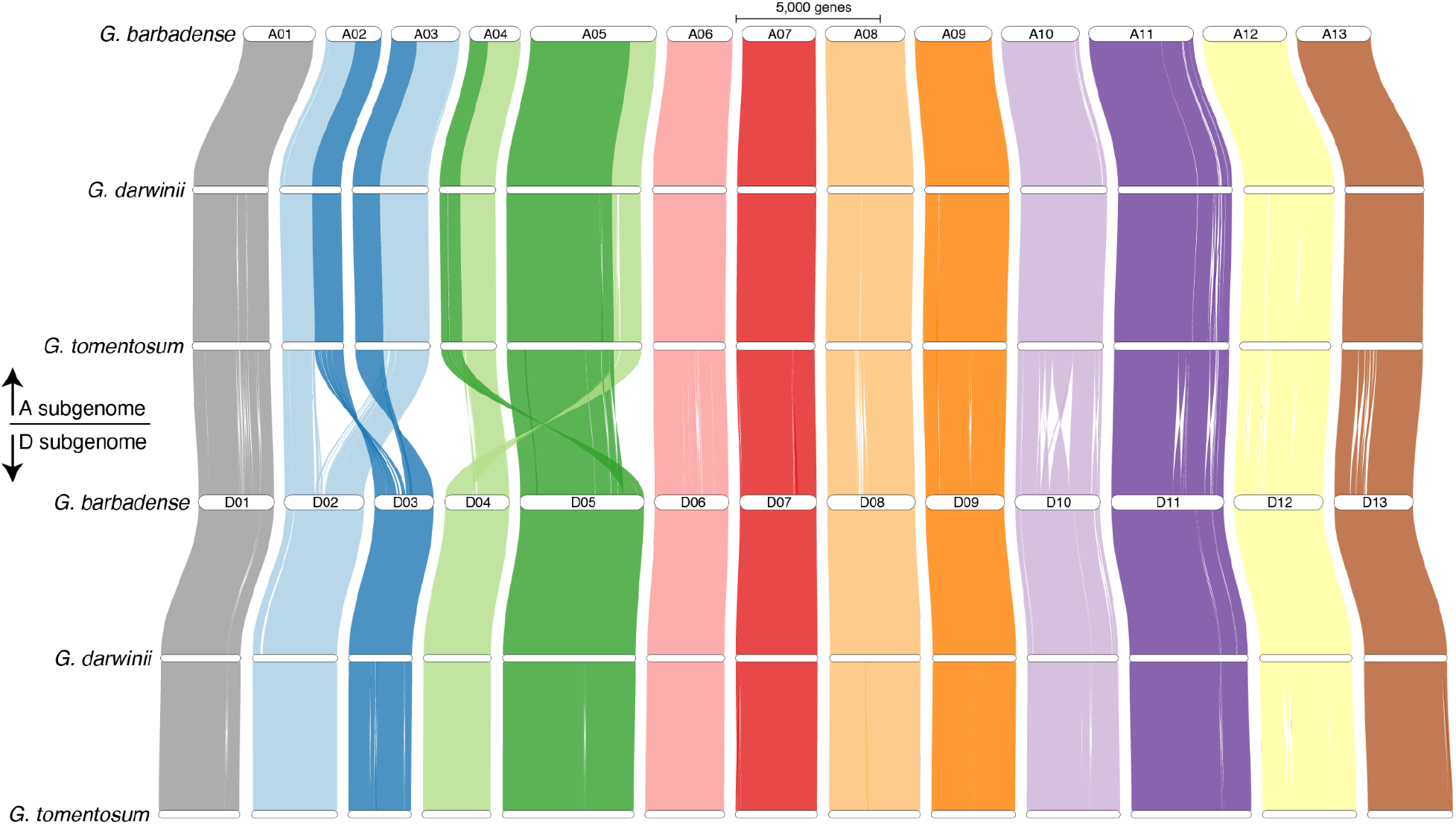
Cotton sub-genome synteny. The synteny map for the split-sub-genome run is presented here. The two *G. barbadense* sub-genome chromosomes are labeled; the top three A sub-genome and bottom three D sub-genome chromosomes map to these. Synteny braids are colored following the D sub-genome chromosome order.

**Supplemental Figure 3.**
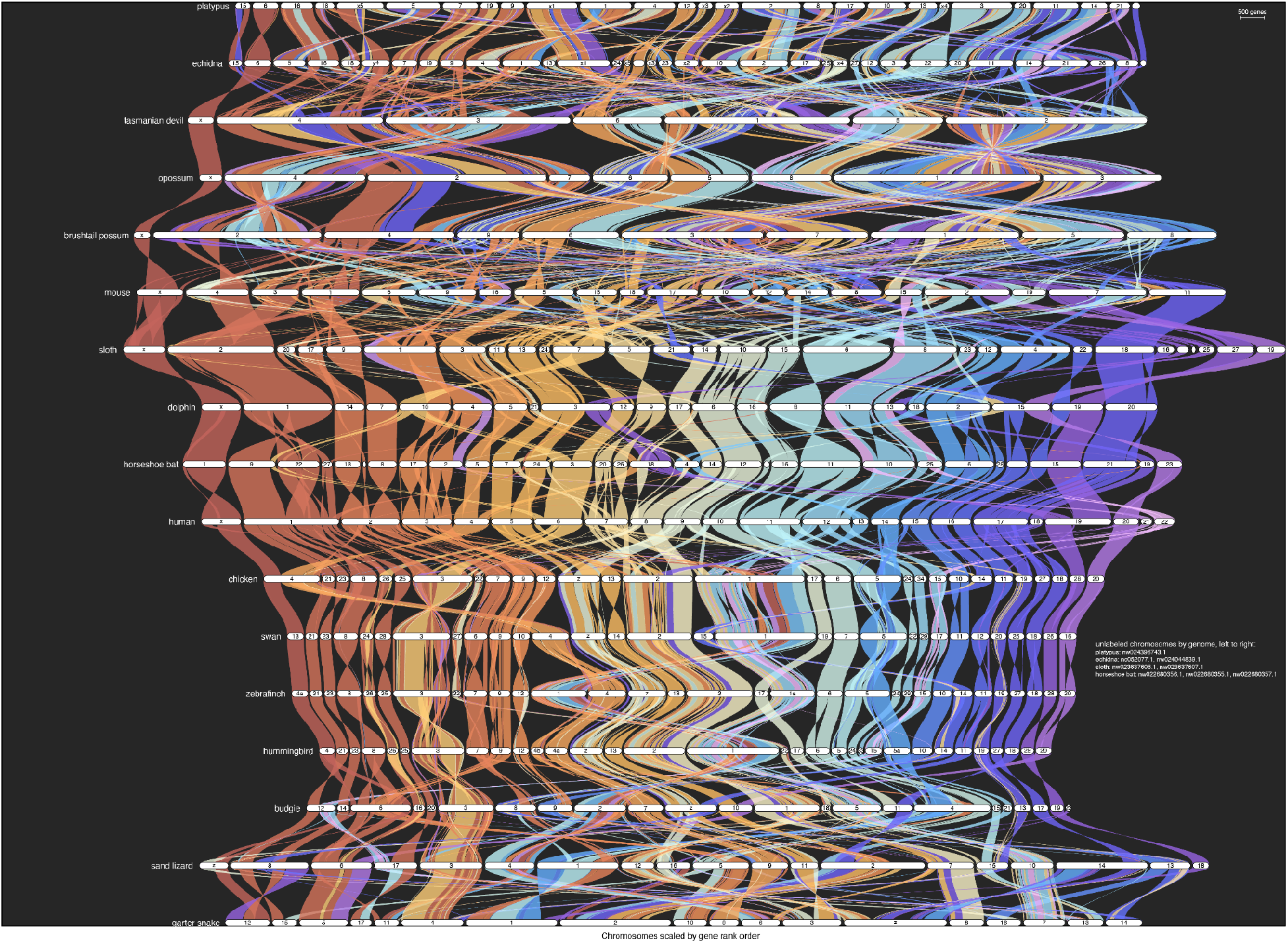
The full synteny map across 17 vertebrate genomes. Chromosomes are ordered to maximize synteny with human chromosomes [X, Y, 1-22]. Syntenic braids are color coded by their mapping to the human chromosomes. A few scaffolds were too small for an informative label. These are listed on the right. Chromosome sizes are scaled by the number of genes with syntenic mappings to other genomes.

**Supplemental Figure 4.**
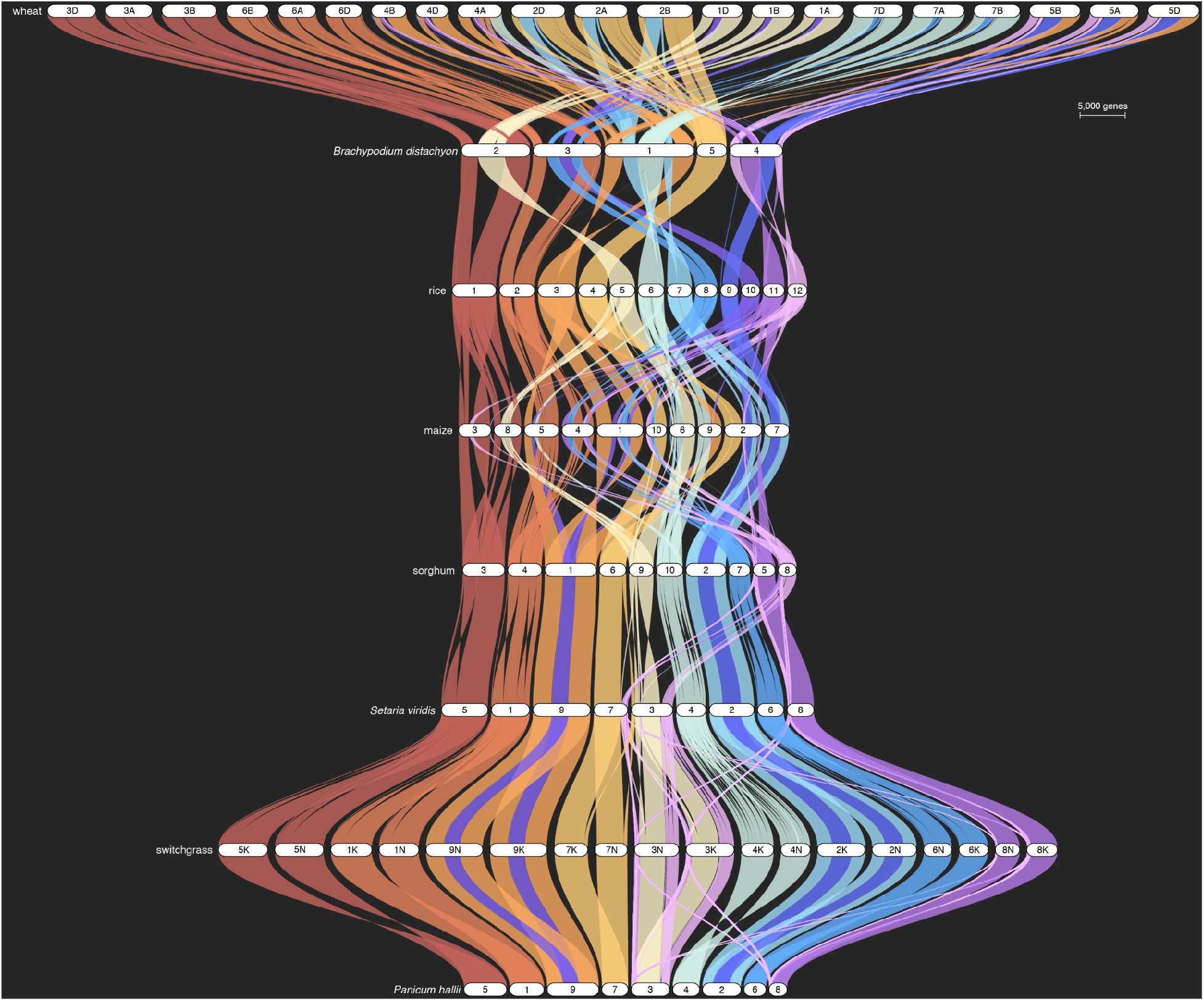
The full synteny map across 8 grass genomes. Chromosomes are ordered to maximize synteny with rice chromosomes [1-12]. Syntenic braids are color coded by their mapping to the rice chromosomes. Chromosome sizes are scaled by the number of genes with syntenic mappings to other genomes.

**Supplemental Figure 5.**
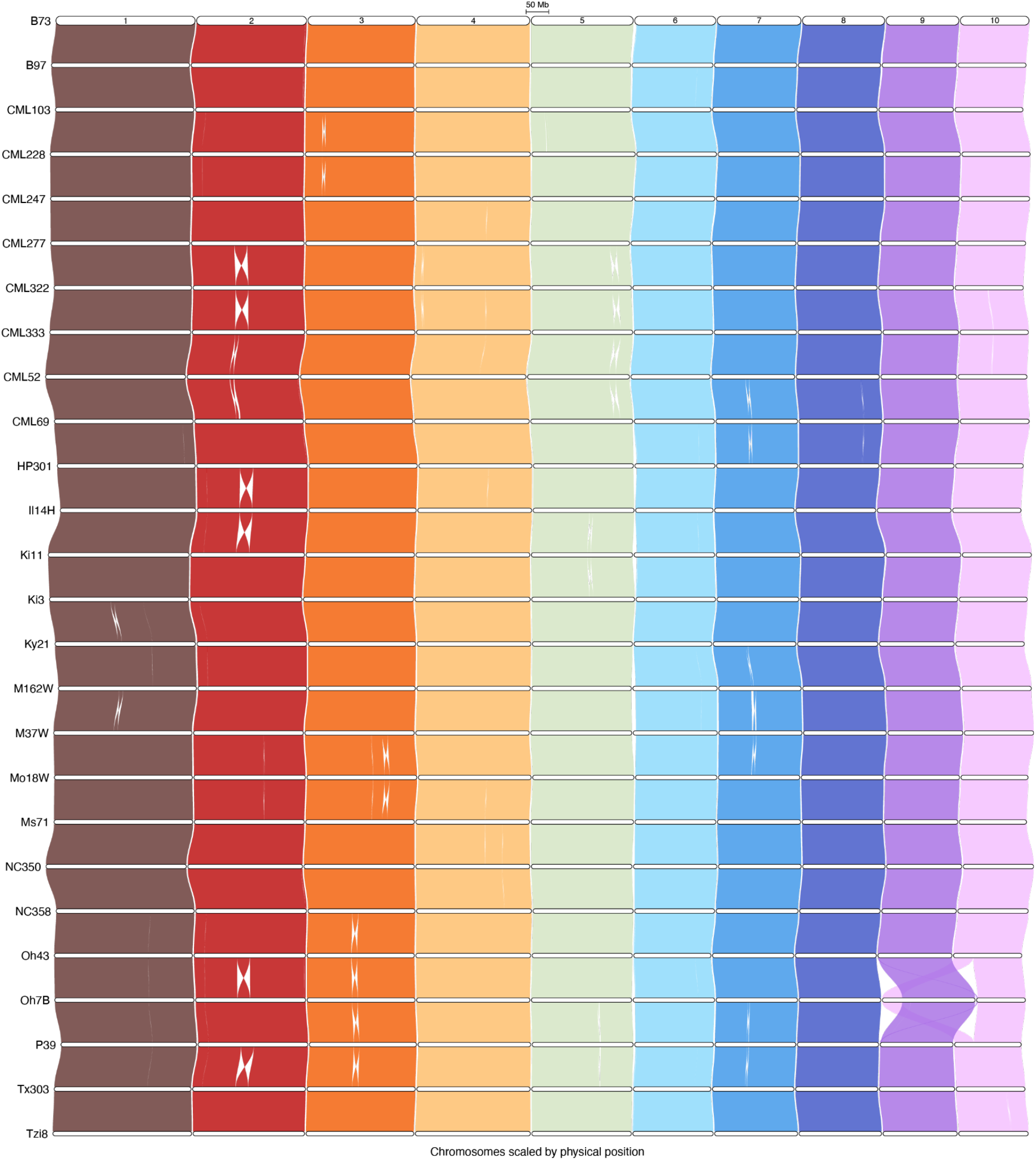
The full synteny map across 26 maize genomes. Chromosomes are ordered 1-10, following the B73 genome labels at the top of the figure. Syntenic braids are color coded by their mapping to the B73 chromosomes. Chromosome sizes are scaled by their physical size.

**Supplemental Figure 6.**
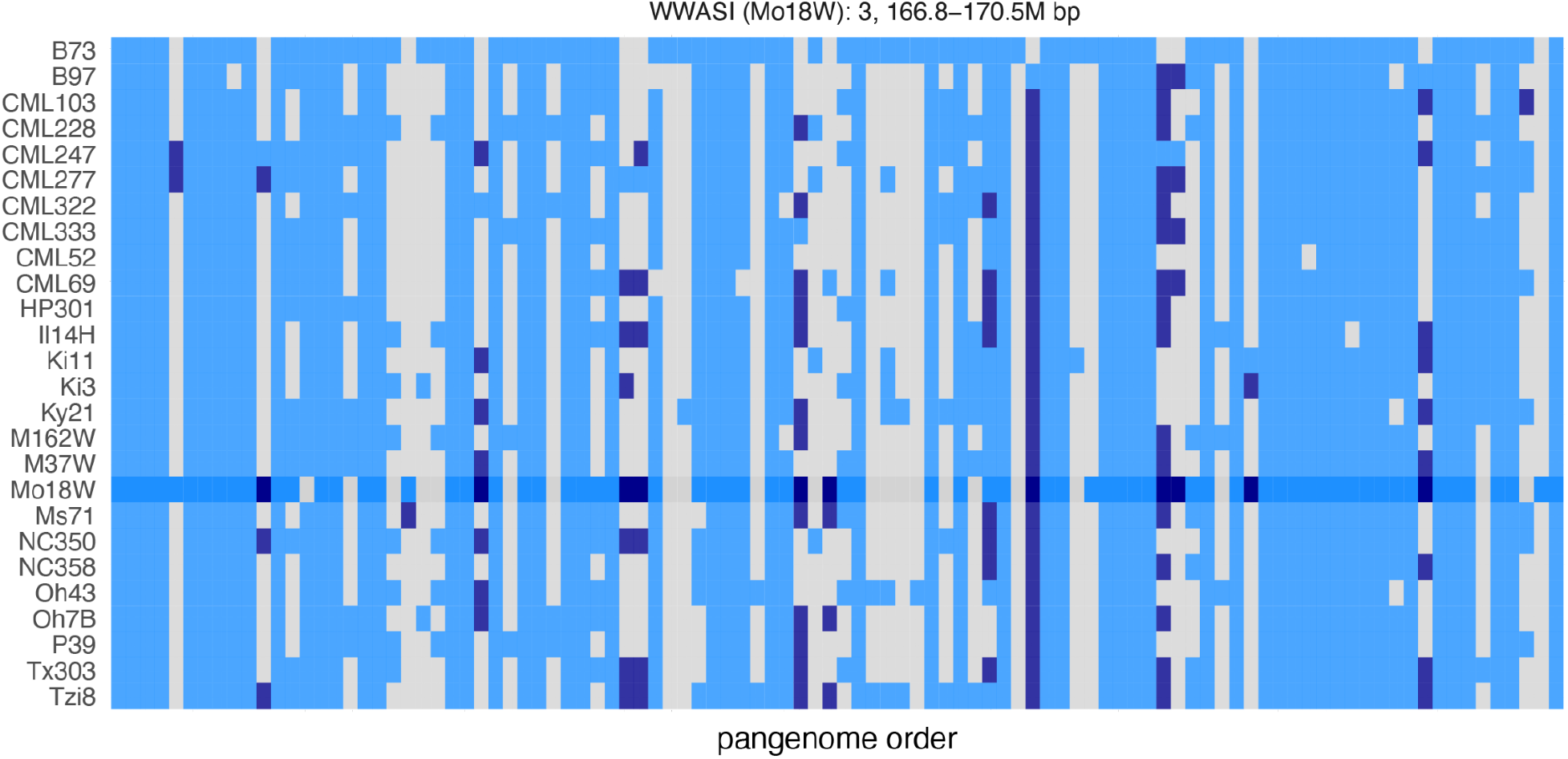
Map of presence absence variation in the larger chromosome 3 QTL interval. Genome labels (y-axis) follow the order of other plots. Pan-genome entries are ordered by physical position within the interval on the x-axis. Gray panes are absences, dark blue are multi-copy and light blue are single-copy genes in each entry-by-genome combination. The more saturated colors correspond to the Mo18W genome, which has an outlier effect on this interval.

